# Whole Genome Bisulfite Sequencing of Cell Free DNA and its Cellular Contributors Uncovers Placenta Hypomethylated Domains

**DOI:** 10.1101/004101

**Authors:** Taylor J. Jensen, Sung K. Kim, Zhanyang Zhu, Christine Chin, Claudia Gebhard, Tim Lu, Cosmin Deciu, Dirk van den Boom, Mathias Ehrich

**Affiliations:** Sequenom Center for Molecular Medicine, 3595 John Hopkins Ct, San Diego, CA, 92121; Sequenom Inc., 3595 John Hopkins Ct, San Diego, CA, 92121

**Author notes:** Email addresses: TJJ SKK ZZ CC CG TL CD DvdB ME. Corresponding author: Taylor Jensen, PhD Group Leader, Research and Development; cell.

**Keywords:** whole genome bisulfite sequencing, DNA methylation, non-invasive testing, circulating cell free DNA, placenta hypomethylation

## Abstract

**Background:** Circulating cell free fetal DNA has enabled non-invasive prenatal fetal aneuploidy testing without direct discrimination of the genetically distinct maternal and fetal DNA. Current testing may be improved by specifically enriching the sample material for fetal DNA. DNA methylation may allow for such a separation of DNA and thus support additional clinical opportunities; however, this depends on knowledge of the methylomes of ccf DNA and its cellular contributors.

**Results:** Whole genome bisulfite sequencing was performed on a set of unmatched samples including ccf DNA from 8 non-pregnant (NP) and 7 pregnant female donors and genomic DNA from 7 maternal buffy coat and 5 placenta samples. We found CpG cytosines within longer fragments were more likely to be methylated, linking DNA methylation and fragment size in ccf DNA. Comparison of the methylomes of placenta and NP ccf DNA revealed many of the 51,259 identified differentially methylated regions (DMRs) were located in domains exhibiting consistent placenta hypomethylation across millions of consecutive bases, regions we termed placenta hypomethylated domains (PHDs). We found PHDs were consistently located within regions exhibiting low CpG and gene density. DMRs identified when comparing placenta to NP ccf DNA were recapitulated in pregnant ccf DNA, confirming the ability to detect differential methylation in ccf DNA mixtures.

**Conclusions:** We generated methylome maps for four sample types at single base resolution, identified a link between DNA methylation and fragment length in ccf DNA, identified DMRs between sample groups, and uncovered the presence of megabase-size placenta hypomethylated domains. Furthermore, we anticipate these results to provide a foundation to which future studies using discriminatory DNA methylation may be compared.

## Background

The field of non-invasive prenatal testing was enabled by the discovery that circulating cell free (ccf) fetal DNA is present pregnant female plasma[1]. It does, however, only present the minority species of DNA in total ccf DNA obtained from pregnant women. This mixture consists of DNA inherently present in the plasma of non-pregnant females, thought to primarily be derived from maternal haematopoietic cells, supplemented with a minority fraction of fetal DNA coming from the placenta [2–5]. Since these nucleic acids are distinct, they can be differentiated through a number of genomic markers including single nucleotide changes, haplotypes, or copy number variants. In addition, DNA methylation can serve as a distinguishing feature and has thus been evaluated for fetal DNA enrichment [6–9]; however, complete analysis requires an in depth knowledge of the genome-wide DNA methylation patterns in ccf DNA isolated from pregnant plasma as well as its primary non-cellular and cellular contributors.

DNA methylation participates in numerous developmental processes including X chromosome inactivation, genomic imprinting, and cellular differentiation [10–13]. Differences in DNA methylation patterns are cell type specific and, in concert with histone tail modifications and other epigenetic alterations, cooperate to modulate chromatin structure [14–17]. While the majority of previous epigenetic studies have been performed upon only a portion of the genome [6, 14, 18–20], recent research from the ENCODE project indicates that up to 80% of the human genome may be functional, highlighting the importance of measuring the DNA methylome in its entirety [21]. Utilizing sequencing techniques that permit complete methylome analysis, a number of studies have described genome-wide methylation profiles of normal and cancer samples [22–30]; however, high resolution methylation maps of complex biological specimens including ccf DNA only recently been described [31].

We performed whole genome bisulfite sequencing (WGBS) [22, 25–27] to characterize the methylome of ccf DNA from eight non-pregnant and seven pregnant female donors. In addition, seven genomic DNA samples isolated from maternal buffy coat and five placenta samples were sequenced at single base resolution. This produced DNA methylome maps for each sample type. The present study provides single-base resolution methylomes of ccf DNA, demonstrates a link between local DNA methylation levels and ccf DNA fragment size, and shows large, continuous regions of hypomethylation in the placenta (Placenta Hypomethlated Domains or PHDs), an epigenetic phenomenon, until recently, only described in tumor samples [24, 30, 32–35].

## Results

Single base resolution methylome maps of ccf DNA isolated from the plasma of 8 non-pregnant female donors were produced using WGBS. We generated 269-551 million paired monoclonal reads per sample, enabling >10x coverage of 74-92% of the ∼28 million genomic CpG sites (Supplementary Figure 1a). Cytosine methylation was evaluated in each of the previously identified genomic contexts (CpG, CHG, and CHH) [26]. Consistent with previous studies on differentiated cell types [36], almost all cytosine methylation occurred in the CpG context with 74.5-75.3% of all CpG cytosines being methylated; methylation in each of the other contexts was minimal (<0.25%; Supplementary Figure 1b). These data created eight comprehensive genome-wide CpG cytosine methylation maps of ccf DNA which can serve as a foundation for subsequent comparisons within this study and beyond (Supplementary Figure 1c).

**Figure 1.**
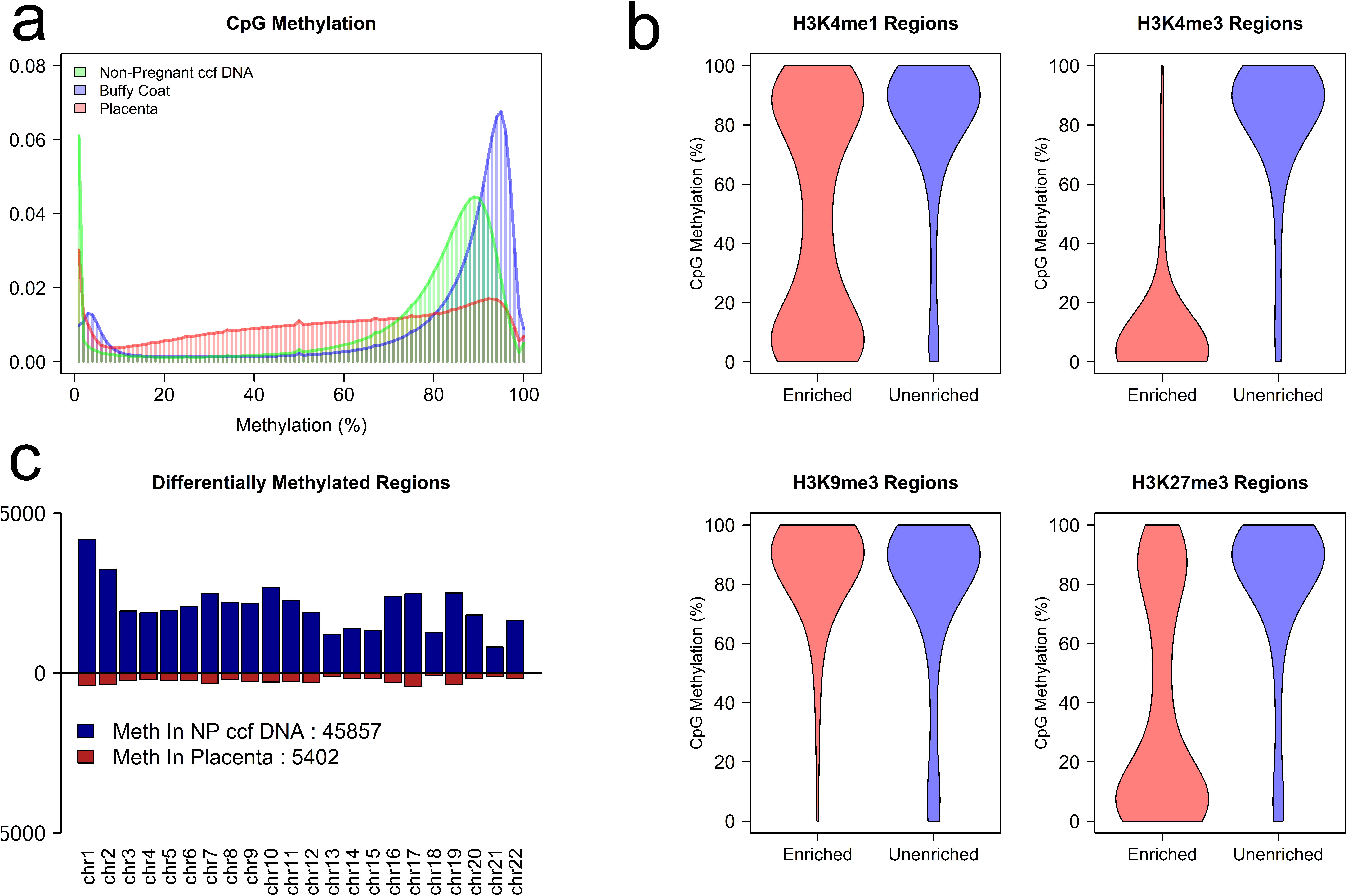
Methylation patterns in buffy coat, placenta, and non-pregnant ccf DNA. a) The distribution of mean CpG methylation for each sample type (nonpregnant ccf DNA, maternal buffy coat, and placenta). The y-axis represents the relative proportion of all evaluated CpG dinucleotides exhibiting a particular level of CpG methylation. The histogram bins each have a width of 1%. b) CpG methylation of non-pregnant ccf DNA samples was assessed in ENCODE-defined enriched regions for H3K4me1, H3K4me3, H3K9me3, and H3K27me3. Unenriched data were generated by a random sampling of the same number of CpG sites as used for enrichment, but located elsewhere in the genome. The width of each violin plot is representative of data density at a given CpG methylation level. c) The number of DMRs more methylated in placenta (red) and non-pregnant (NP) ccf DNA (blue).

Previous work has indicated that the predominant contributor to non-pregnant ccf DNA are cells of hematopoietic origin [4]. This led us to perform WGBS on buffy coat cells obtained from 7 distinct pregnant female donors (Supplementary Figure 2). Methylation levels at 37775 CpG sites were confirmed by MassARRAY in an independent sample cohort of 8 buffy coat samples (Pearson correlation=0.953; Supplementary Figure 3). Nearly all CpG sites in buffy coat showed either low (9.7%; defined as less than 20% mean methylation across all buffy coat samples) or high (79.8%; defined as greater than 75% mean methylation across all buffy coat samples) levels of methylation (Figure 1a), similar to the distribution in non-pregnant ccf DNA.

**Figure 2.**
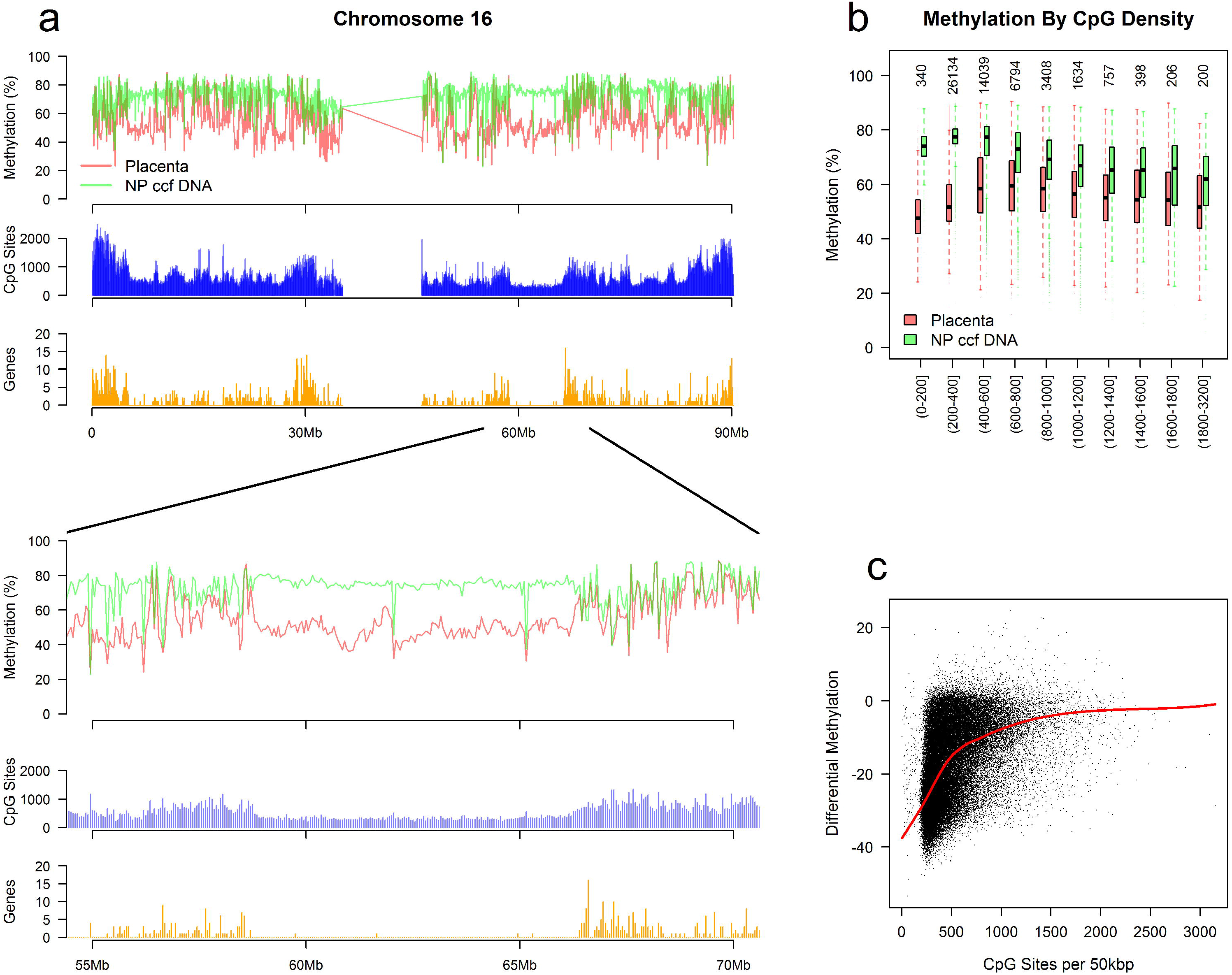
Identification of Placenta Hypomethylated Domains (PHDs). a) Mean methylation per 50 kbp genomic bin on chromosome 16 with non-pregnant ccf DNA (NP ccf DNA) and placenta shown. CpG sites (blue) and genes (orange) were summed per 50kbp genomic bin. b) Genomic methylation level by CpG density at 50kbp bin level. Values on the x-axis represent the number of CpG sites per 50kbp bin. Numbers along the top indicate the number of genomic bins analyzed. c) Differential methylation between placenta and non-pregnant plasma as a function of CpG density at 50kbp bin level. A negative value on the y-axis is indicative of placenta hypomethylation. The red line corresponds to a loess smoothed fit.

**Figure 3.**
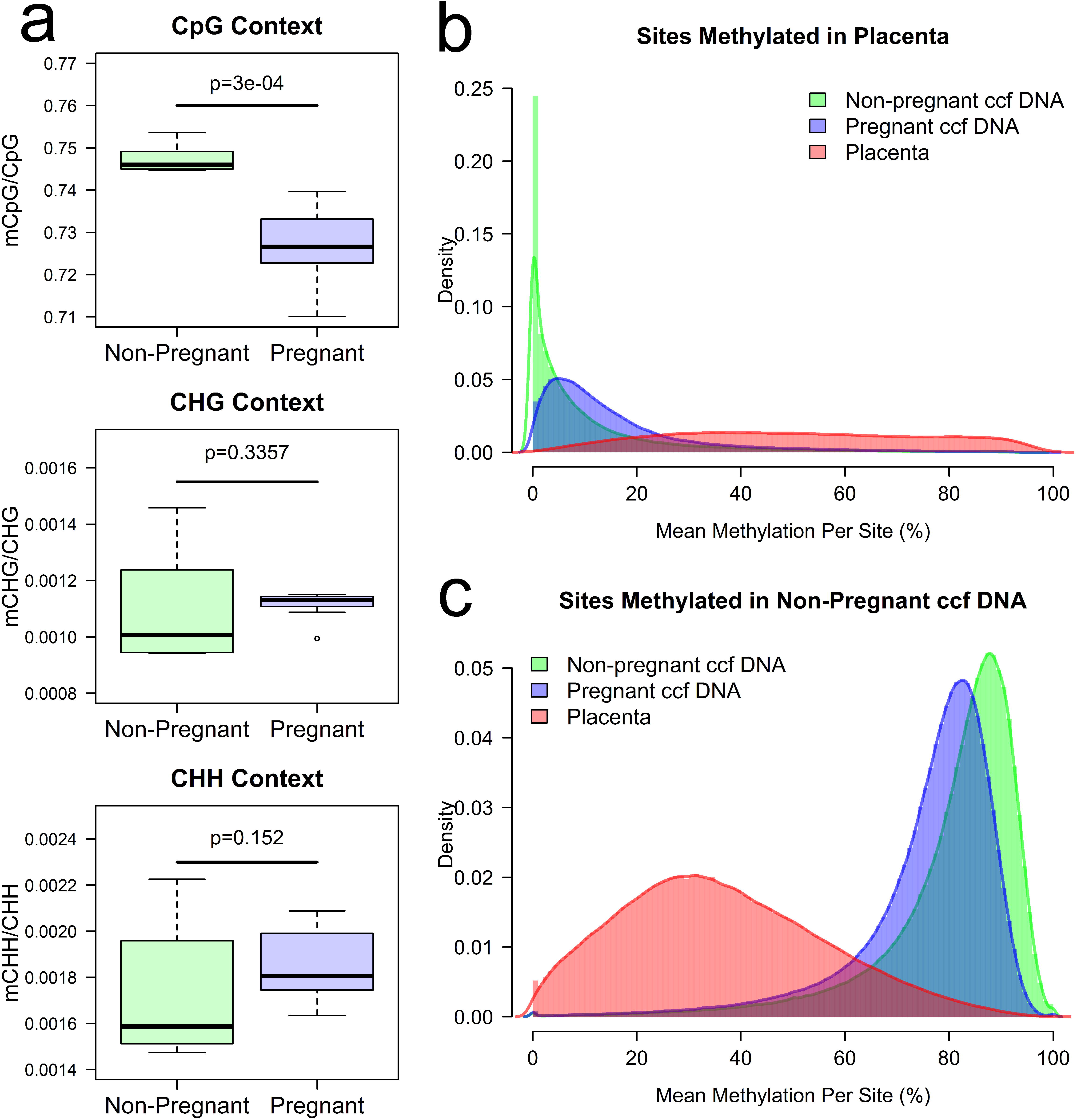
Methylome of ccf DNA isolated from pregnant plasma. a) Cytosine methylation in non-pregnant and pregnant ccf DNA for CpG, CHG, and CHH contexts are shown. P-values were calculated using a Wilcox rank sum test. b) Methylation of all cytosines located within the DMRs hypermethylated in placenta tissue relative to non-pregnant ccf DNA. The y-axis (density) is the defined as the proportion of CpG sites at a given methylation level. c) Methylation of all cytosines located within the DMRs hypermethylated in non-pregnant ccf DNA relative to placenta tissue. The y-axis (density) is the defined as the proportion of CpG sites at a given methylation level.

Next, the link between histone tail modifications and DNA methylation was examined. Publically available PBMC ChIP-Seq data from the ENCODE project were used to identify regions enriched for four distinct histone H3 modifications. Since the methylome of non-pregnant ccf DNA closely resembled that of buffy coat (PBMC), the level of CpG methylation in non-pregnant ccf DNA was then examined within these regions. (Figure 1b). In regions enriched for H3K4me3, 89.9 % of cytosines showed less than 20% methylation while only 5.2% of unenriched sites were similarly unmethylated. Conversely, 84.9% of CpG sites were methylated (>75%) in H3K9me3 enriched regions as compared to 76.3% in unenriched regions. Distinct differences were also observed when comparing H3K4me1 and H3K27me3 enriched regions to corresponding unenriched CpG sites. Taken together, these data suggest a link between particular histone marks and CpG methylation in buffy coat. Comparison of the methylomes of buffy coat and non-pregnant ccf DNA indicated high similarity (Pearson correlation=0.954; Supplementary Figure 4); however, we detected 152 differentially methylated regions (DMRs) (139 more methylated in buffy coat), suggesting there are additional sources of cell free DNA distinct from buffy coat present in circulation. These data link histone modifications to CpG methylation in buffy coat and suggest that the majority of ccf DNA is derived from the hematopoetic compartment with minimal contributions from alternative tissues.

**Figure 4.**
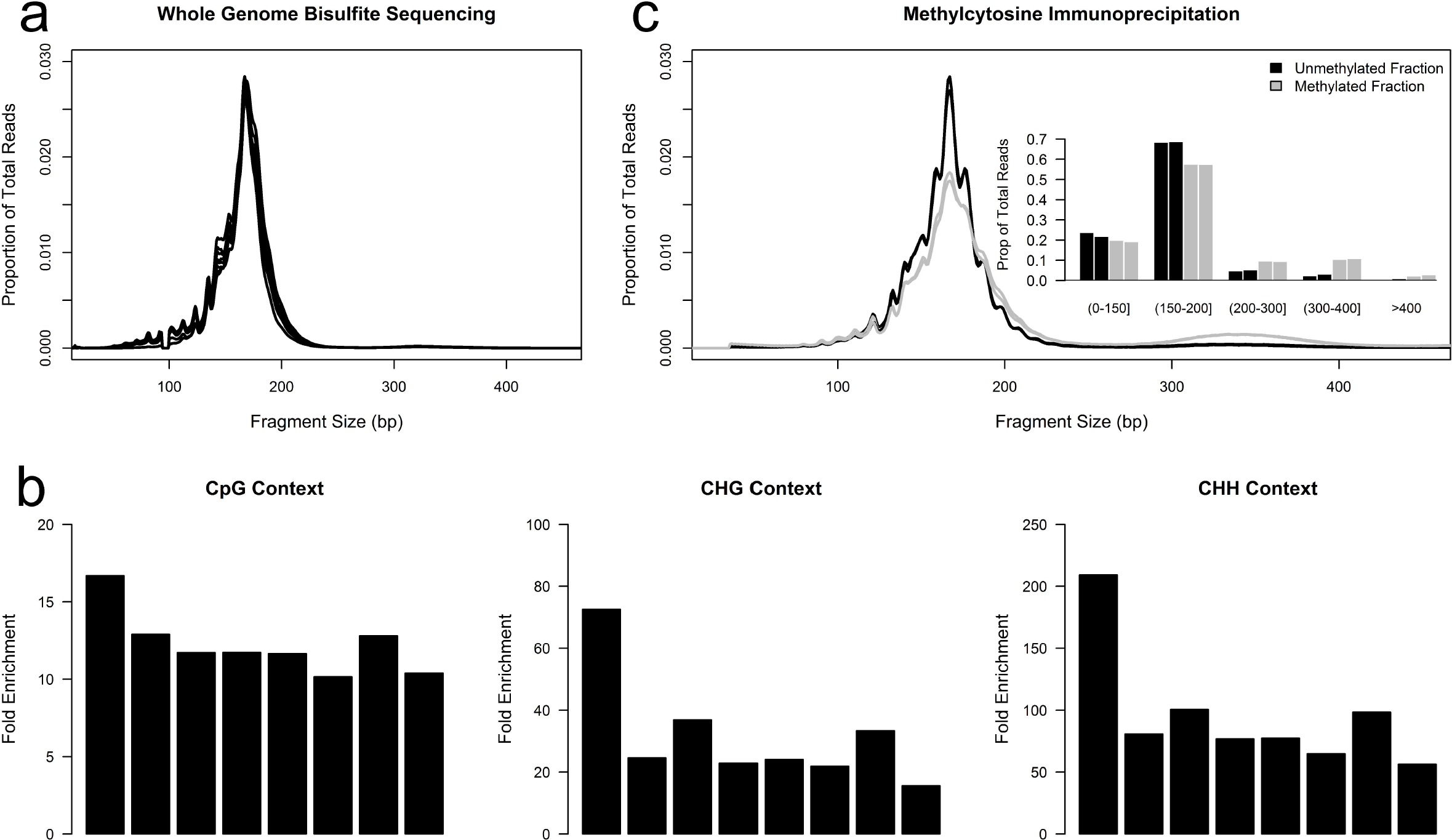
Linkage between fragment size and local DNA methylation in nonpregnant ccf DNA. a) Fragment size of ccf DNA as measured by WGBS. Each line represents an individual ccf sample. Loss of representation at ∼92-98 bp is an artifact of adapter trimming prior to alignment. b) Ratio of methylated CpG, CHG, and CHH cytosines within large fragments (>200bp) relative to methylated cytosines in small fragments (<200bp) after scaling for number of cytosines measured. Each bar represents a single sample. c) Size distribution of unmethylated fraction (black) and methylated fraction (gray) using MCIp-Seq. Each line represents an individual sample fraction. Inset plot provides a quantified distribution of fragment size for each fraction.

Since the fetal portion of ccf DNA in pregnant plasma is derived from the placenta [2–5], WGBS of 5 placenta samples was performed to identify of placenta specific DMRs (Supplementary Figure 2). Methylation levels of 37,775 CpG sites were also measured using MassARRAY in a separate eight sample cohort and showed high concordance (Pearson correlation=0.897; Supplementary Figure 5). Comparison of the distribution of methylation in placenta to the distribution in non-pregnant ccf DNA or buffy coat revealed a significant difference (p<2.2e-16; Kolmogorov-Smirnov Test). While only 15.5% and 10.5% of CpG sites exhibited intermediate methylation (20%-75% mean methylation) in non-pregnant ccf DNA and buffy coat, respectively, 46.6% of CpG sites showed intermediate methylation in placenta tissue (Figure 1a). Comparison of CpG sites between placenta and buffy coat revealed that the majority of the intermediate methylated regions in placenta were highly methylated in both non-pregnant ccf DNA and buffy coat (Supplementary Figures 6,7). CpG methylation was compared to gene expression determined by microarray analysis on an independent cohort of 8 placenta samples. Transcription start sites (TSS) were generally unmethylated independent of gene expression level, while promoter and intragenic regions were linked to gene expression (Supplementary Figure 8).

Differential methylation between placenta and each of the aforementioned sample types was then analyzed. We identified 51,259 DMRs between placenta and nonpregnant ccf DNA, of which 89.5% were more methylated in ccf DNA, consistent with the observed distribution differences (Figure 1c; Supplementary Figure 9). We assayed 243 of the putative DMRs in an independent sample set of 6 placenta samples and 8 non-pregnant ccf DNA samples using MassARRAY and 98.8% (240/243; Supplementary Figure 10) were confirmed (p<0.05; Wilcox Rank Sum).

Interestingly, these DMRs overlapped with CpG islands in only 7.9% of cases and frequently occurred within intragenic and intergenic regions (Supplementary Figure 11). This may be due to the observed low CpG methylation levels within CpG islands in non-pregnant ccf DNA, whereas other genomic regions, including introns and exons, were largely methylated in non-pregnant ccf DNA and hypomethylated in placenta (Supplemental Figure 12). In addition, we identified 105,874 DMRs between placenta and buffy coat with a similar overrepresentation (94.7%) of buffy coat specific methylated regions (Supplementary Figures 13, 14). The majority (93.6%) of DMRs identified between ccf DNA and placenta were also identified as DMRs between placenta and buffy coat (Supplementary Figure 15). Comparison of methylation between buffy coat and placenta in the context of ENCODE defined histone modifications revealed an interesting pattern. Little difference in methylation is observed within H3K4me3 regions while a dramatic difference occurs in H3K9me3 and H3K27me3 enriched regions (Supplementary Figure 16). Regions enriched for H3K4me1 show a generalized decrease in DNA methylation levels in placenta tissue relative to ccf DNA from non-pregnant plasma (Supplemental Figure 16). These differences may indicate differential histone modification profiles within the placenta relative to buffy coat or differences in the correlation between these marks in the placenta. These data provide a genome-wide map of placenta specific DMRs when compared to either non-pregnant ccf DNA or buffy coat.

Examination of the genomic distribution of differential methylation uncovered large contiguous genomic regions with significant placental hypomethylation relative to non-pregnant ccf DNA. We termed these regions PHDs and found that these domains overlapped with a substantial portion (29.9%) of all autosomal hypomethylated DMRs. PHDs were characterized by a number of distinguishing characteristics.

First, they were typically located in regions of low CpG and gene density (gene deserts). Secondly, these regions often exhibited high levels of DNA methylation in ccf DNA from non-pregnant plasma (mean methylation 74.2%) while placenta tissue showed a considerably lower level of methylation (mean methylation 49.9%). Using a window size of 50kbp, we detected PHDs on each autosome that covered as many as ∼14 million bases. Figure 2a shows a number of these regions located on chromosome 16 with particular focus upon a 7.5 Mbp PHD located on chromosome 16q. Since the presence of a PHD was consistently observed in regions of low CpG density, the link between CpG density and methylation levels was further examined. Indeed, the magnitude of placenta hypomethylation in low CpG density regions far surpasses that observed in more dense regions (Figure 2b). A similar pattern is seen when comparing CpG methylation to gene density (Supplementary Figure 17). Moreover, the magnitude of differential methylation was positively linked to the local CpG density (Figure 2c). These data identify large genomic regions which are consistently hypomethylated in the placenta and link these regions to low CpG and gene density. While additional work is needed to further characterize PHDs, these characteristics perhaps underscore a lack of heterochromatin formation during early placenta development or allele specific methylation of regions with low CpG density in the placenta [37].

We measured the methylome of ccf DNA derived from the plasma of seven pregnant female donors to see if we could detect the DMRs identified between placenta and non-pregnant ccf DNA (Supplementary Figure 2). Overall methylation levels in pregnant and non-pregnant ccf DNA were similar for non-CpG cytosines (<0.25%); however, overall methylation within a CpG context was significantly reduced from 74.5-75.3% to 71.0-74.0% (p=3e-04, Wilcoxon rank-sum; Figure 3a). Since ccf DNA from pregnant plasma is comprised of maternal and fetal ccf DNA, methylation patterns should be a composite of non-pregnant ccf DNA and placenta tissue. To address this, we evaluated the mean methylation level of each CpG site within DMRs identified between non-pregnant ccf DNA and placenta. CpG sites within identified DMRs exhibited significantly (p<2e-16; Wilcoxon rank-sum) different methylation levels in pregnant ccf DNA relative to non-pregnant ccf DNA (Figures 3b, 3c; Supplementary Figures 18, 19). Hierarchical clustering confirmed these results by clustering pregnant and non-pregnant ccf DNA samples as single branches on a dendrogram (Supplementary Figure 20,). Overall, these data confirm the differential methylation identified when comparing non-pregnant ccf DNA and placenta tissue.

Previous reports have indicated that fetal ccf DNA is shorter than its maternal counterpart [38–40]. Since hypomethylation is linked to an open chromatin structure and thus may exhibit an increased accessibility to native endonucleases during apoptosis [41], we assessed the relationship between CpG methylation and ccf DNA length in non-pregnant plasma to determine if this contributes to the observed size difference. In each of the samples analyzed, the most prominent length was 168bp, similar to previous reports (Figure 4a) [38]. After accounting for the differences in the number of analyzed bases for each size fraction, we found that CpG cytosines within longer fragments (>200bp) were on average 12.3-fold more likely to be methylated (Figure 4b). Interestingly, a similar pattern was also found for cytosines in the CHG (31.5-fold) and CHH (95.5-fold) contexts, although their overall occurrence was much lower than methylated CpG cytosines. A similar relationship between CpG methylation likelihood and fragment length was also observed in ccf DNA from the plasma of pregnant women but was not observed in the manually sheared buffy coat and placenta samples (Supplementary Fig 21), consistent with this relationship being the result of biological DNA fragmentation. We performed methyl-CpG immunoprecipitation (MCIp)-Seq on an independent set of two nonpregnant ccf DNA samples to confirm the observed size differences for CpG cytosines. MCIp enables the separation and collection of both the unmethylated and methylated fractions of a sample. Sequencing both fractions from each sample revealed a distinct size difference with the most striking difference between fractions occurring at ∼320 bp, roughly the size of two nucleosomes (Figure 4c). Indeed, the proportion of DNA fragments greater than 300bp is 3.8-fold higher in methylated fragments (13.3%) than in unmethylated fragments (3.5%; Figure 4c). Conversely, the proportion of short (<100bp) ccf DNA fragments is increased in regulatory regions including promoters (4.8%) and CpG islands (8.2%) relative to the entire genome (2.2%; Supplemental Figure 22). Overall, these data link DNA methylation and potentially other epigenetic marks to fragment length in ccf DNA.

Non-invasive prenatal aneuploidy detection is linked to the fraction of fetal (placental) DNA in the sample [42–45]. We hypothesized that the global hypomethylation of the placenta may allow enrichment for fetal DNA. We isolated ccf DNA from the plasma of an independent set of 12 pregnant donors, three of which were confirmed to be carrying a fetus affected with trisomy 21, and measured each sample with and without enriching for unmethylated DNA. Data from a subset of PHDs showed that enriching for unmethylated DNA resulted in a 3.99-fold (range: 2.9-5.9 fold) increase in chromosome 21 z-scores in trisomy 21 samples relative to the same samples without enrichment; one sample from a euploid pregnancy showed a similar level of enrichment (Supplementary Figure 23). Overall, while the sample size is small, these data suggest that placenta hypomethylation may be leveraged to increase the effective fetal fraction in pregnant ccf DNA samples.

## Discussion

We created whole genome methylome maps for a total of 27 samples from 4 distinct sample types, enabling a comprehensive characterization of the methylome of ccf DNA from pregnant plasma and each of its primary cellular and non-cellular contributors. We identified a total of 152 DMRs when comparing non-pregnant ccf DNA to DNA isolated from buffy coat, thought to be the primary cellular contributor to this nucleic acid pool. While the DNA methylation patterns are similar (Pearson correlation=0.954), the differences identified are consistent with additional minority contributors to non-pregnant ccf DNA. Further studies are required to determine the identity of additional contributors, but sources may include organ systems with extensive bloodstream contact including the kidneys, liver, or endothelium. We also identified 51,259 DMRs when comparing placenta to non-pregnant ccf DNA. Previous studies have identified placenta specific methylated sites within subsets of the genome [18–20]. In each of these studies, the authors have used a model system consisting of placenta and buffy coat/PBMC to identify these DMRs. Results from our study show a much greater number of differentially methylated regions when comparing buffy coat to placenta (105,874), suggesting a higher false positive rate when using this genomic DNA model system alone.

While we identified genome-wide placental hypermethylated regions consistent with previous studies, we have also leveraged the global hypomethylation patterns in the the placenta for an initial proof of concept study for fetal enrichment. Specifically, we evaluated the principle of global hypomethylation as a method of enriching for fetal DNA in a set of 12 ccf DNA samples from pregnant female donors, three of which carried a fetus with trisomy 21 (T21). Using a z-score cutoff of three to suggest an overrepresentation of chromosome 21 in the samples enriched for unmethylated DNA, all three of the T21 samples were detected. In addition, there was one euploid sample which exhibited similar enrichment and thus would be categorized as a false positive using this classification criteria. While these data are promising as an early proof of concept, further work is needed to evaluate the robust performance of DNA hypomethylation as a method for fetal DNA enrichment in ccf DNA derived from the plasma of pregnant donors.

This study was designed to evaluate the proposed major contributors of nucleic acids into the plasma of a pregnant individual. As part of this design, independent, unpaired samples were used for each of the discovery and confirmatory processes. While using a paired study design would have improved the continuity of the comparisons between methods, we hypothesized that this unpaired study design would produce a higher likelihood that the results are reproducible across a larger sample set. Furthermore, since the methylation patterns in ccf DNA from pregnant plasma were consistent with the regions we identified in placenta samples despite differences in gestational age between these sample types (Supplementary Table 1), the identified differences are likely stable during early gestation; however, since all ccf DNA and placenta samples were obtained from donors at less than 25 weeks, we cannot rule out that changes in DNA methylation occur within these regions during late gestation.

While evaluating the genomic distribution of DMRs, we unexpectedly observed large regions of placental hypomethylation. These data are reinforced by a recent study which identified a similar pattern in a single placenta sample using low coverage WGBS [35]. Further characterization of these regions indicated that they were present in regions with low CpG and gene density. Regions with these characteristics are often located within heterochromatinized domains, pointing to a reduction in the formation or re-distribution of heterochromatin in the developing placenta. This is supported by the observed decrease in CpG methylation in the placenta within regions containing the H3K9me3 mark in PBMC (Supplemental Figure 15). The identified PHDs showed characteristics consistent with the partially methylated domains and/or global hypomethylation previously described in cancer subtypes [24, 30, 33]. Commonalities between the placenta and tumors have been previously described and include an increased proliferation rate, the ability to migrate, and invasive potential [46]. These data indicate that the parallels between cancer and the placenta extend to their epigenomes and may provide an experimental opportunity for elucidating the molecular source of these similarities. In addition, such similarities suggest that lessons learned from this study may be directly applicable to non-invasive tumor detection and monitoring.

## Conclusions

This project enabled the generation of methylome maps for each sample type at single base resolution, identified a link between local DNA methylation and fragment length of ccf DNA, provided comprehensive lists of differentially methylated regions (DMRs) between sample groups, and uncovered the presence of megabase-size PHDs. Taken together, this study advances the biological understanding of ccf DNA and placenta. Furthermore it delivers the ccf DNA methylome at single base resolution as a reference for future non-invasive diagnostic studies.

## Methods

### Blood processing and DNA extraction

Plasma samples were collected under two separate Investigational Review Board (IRB) approved clinical protocols (BioMed IRB 301-01 and Western IRB 20090444). Buffy coat and placenta tissue was collected from consented subjects under a Western IRB approved protocol (20111833, study #1128724) and in accordance with the FDA Guidance on Informed Consent for in vitro Diagnostic Device Studies Using Leftover Human Specimens that are Not Individually Identifiable (April 25, 2006). All subjects provided written informed consent prior to undergoing any study related procedures. All information was anonymized prior to processing. Blood was processed and DNA extracted as previously described [42, 44, 47]. Further information about all samples subjected to WGBS is supplied in Supplemental Table 1.

### Library preparation of ccf DNA

For libraries created from ccf DNA, DNA was subjected to end repair, mono-adenylation, and ligation as previously described[43, 47]. Ccf DNA exists as small fragments and thus no size selection is required prior to sequencing; therefore, the length of each library insert reflects of native DNA fragment length. Ligated products were treated with sodium bisulfite (EpiTect; Qiagen) using a cycling incubation of 95°C for 5 minutes, 60°C for 25 minutes, 95°C for 5 minutes, 60°C for 85 minutes, 95°C for 5 minutes, and 60°C for 175 minutes followed by 3 cycles of 95°C for 5 minutes, 60°C for 180 minutes. Each reaction was purified according to the manufacturer’s instructions (Qiagen). Converted product was amplified using Pfu Turbo Cx Hotstart DNA polymerase (Agilent) and the TruSeq primer cocktail (Illumina) using the following cycling parameters: 95°C for 5 minutes; 98°C for 30 seconds; 14 cycles of 98°C for 10 seconds, 65°C for 30 seconds, 72°C for 30 seconds; and 95°C for 5 minutes.

### Library preparation of genomic DNA

For libraries created from buffy coat or placenta tissue, genomic DNA (10 μg) was fragmented by sonication and column purified (Qiagen). Three ligated products were prepared from each sample (2.5 μg each) by performing end repair, mono-adenylation, and adapter ligation according to the manufacturer’s protocol (TruSeq; Illumina). Bead-based purification (AMPure XP; Beckman Coulter) was performed after the end repair and ligation processes. Ligated products were pooled and 2 distinct bisulfite conversion reactions were performed as described above. Eluted products from each sample were pooled and concentrated using a column-based method (Qiagen). Finally, 40% of each converted sample was amplified as described above. PCR products were purified using magnetic beads (AMPure XP; Beckman Coulter).

### Methyl-CpG immunoprecipitation (MCIp) library preparation

Ccf DNA was isolated from the plasma of either two non-pregnant female donors or twelve pregnant female donors and subjected to methyl-CpG immunoprecipitation according to the manufacturer’s instructions (EpiMark; New England Biolabs).

Briefly, DNA was incubated with the MBD-Fc protein in the presence of 150mM NaCl. DNA which did not bind to the protein was collected and characterized as the unmethylated fraction. The protein-DNA complex was washed three times with 150 mM NaCl and DNA was eluted by heating to 65°C for 15 minutes. Resultant unmethylated and methylated fractions from each donor sample were subjected to library preparation using a modified version of the manufacturer’s protocol. Due to low input amounts, adapter ligation was performed using a diluted adapter oligonucleotide (1:10 for unmethylated; 1:100 for methylated). Resultant ligated ccf DNA was amplified using TruSeq PCR Master Mix and TruSeq primer cocktail (Illumina) using the following cycling parameters: 98°C for 30 seconds; 10 cycles of 98°C for 10 seconds, 65°C for 30 seconds, 72°C for 30 seconds; and 72°C for 5 minutes.

### Massively parallel sequencing

Library quantification and flowcell clustering were performed as previously described[42, 44, 47]. Paired end sequencing was performed for 100 cycles for all whole genome bisulfite samples and 36 cycles for all MCIp-seq samples.

### Whole genome bisulfite sequencing analysis

Libraries prepared from Phi-X were sequenced upon each flowcell to ensure accurate basecalling. All methylation analysis was performed using v0.9.0 of the Illumina bisulfite sequencing analysis program. Bismark v.06.3[48] was utilized to align each sequenced read to a bisulfite converted human genome (hg19) using Bowtie v.0.12.7[49] and simultaneously perform cytosine methylation calls. Prior to alignment, each read was trimmed to remove contaminating adaptor sequences. Each trimmed sequence read was then aligned to each of four bisulfite converted genomes, each derived from the conversion of each strand and the corresponding complement. Alignment was determined by the single best alignment score to one genome. Methylation was subsequently called for each covered cytosine and summary statistics calculated using the Bismark methylation_extractor script.

### MCIp sequencing analysis

Data were aligned to the February, 2009 build of the human genome (hg19) allowing for only perfect matches within the seed sequence using Bowtie. All paired reads with an insert size greater than 500 bp (0.1-0.4% of all sequencing reads) or with discordant chromosome mapping results were discarded prior to analysis. Size was calculated as the distance between the start site of each of the two paired end reads.

### Post analysis processing

Post analysis processing was performed using custom scripts in an R or perl programming environment. Under the assumption that strand specific methylation is uncommon in ccf DNA, methylation calls mapped to the reverse strand were converted to their corresponding forward strand positions and methylation levels recalculated prior to all analyses. The location of each genomic region was obtained from the hg19 build of the UCSC genome browser. Length of each read was calculated by subtracting the distance of the start position of each paired read. The ENCODE data for the four histone tail modifications in PBMC samples was downloaded as narrowPeak files from the UCSC genome ENCODE site.

### DMR Identification

The mean and standard deviation were calculated for each covered CpG site for each sample type. A t-statistic was then calculated for each CpG site for all comparisons. All sites with a t-statistic with an absolute value less than 5 were removed. CpG sites were grouped if there was less than 300bp between them after t-statistic filtering. A region was then considered a DMR if there were 9 or more CpG sites present.

### EpiTYPER (MassARRAY) Analysis

EpiTYPER analysis was performed as previously described[50]. Samples used for EpiTYPER analysis were distinct from those used for WGBS. To confirm WGBS methylation levels, an independent set of 8 placenta villi samples and 8 maternal buffy coat samples were used. An additional independent set of 6 placenta villi samples and 8 non-pregnant ccf DNA samples were used for DMR validation. Regions were selected for DMR validation using EpiTYPER if they were located on chromosomes which most commonly exhibit trisomies (chromosomes 13, 18, and 21) and if they were hypermethylated in placenta tissue relative to ccf DNA from nonpregnant plasma.

### Gene Expression Analysis

RNA was extracted from an independent set of eight placenta villi samples according to manufacturer’s protocol (Qiagen) and hybridized to Affymetrix Human Exon 1.0 ST microarrays. All raw data files (.CEL) were subjected to rma-sketch normalization using Affymetrix Power Tools scripts. Results were subsequently filtered to remove all transcripts which were not included as part of the main array design (4219) and transcripts without a defined gene (329), leaving a final set of 17,463 genes. All genes without a defined TSS as part of the refseq or Ensembl gene lists or those not located on autosomes were discarded, leaving a final set of 16,231 genes. These genes were subsequently tiered into the high (5,410), low (5,411), and intermediate (5,410) expressing genes.

### MCIp Trisomy Evaluation

Ccf DNA was extracted from two aliquots of plasma (4 mL each) collected from 12 pregnant female donors, three of which were carrying a fetus affected with trisomy 21. The ccf DNA from each sample was then pooled to minimize any collection bias and subsequently separated into two aliquots. Aliquots were then either left untreated or subjected to MCIp to enrich for unmethylated DNA. Sequencing libraries were prepared and sequenced as described above. All data which aligned within a subset of the identified placenta hypomethylated regions were used for downstream analysis. The median and median absolute deviation (MAD) were calculated using data from known euploid samples only for both unenriched and enriched samples independently. Depending on the group (unenriched vs enriched), chromosome 21 z-scores were calculated using a robust method as follows: Z=(Chr 21 Fraction_sample_-Chr 21 Fraction_Median_)/Chr 21 Fraction_MAD_.

## Abbreviations used

ccf: (circulating cell free);
WGBS: (whole genome bisulfite sequencing);
DMRs: (differentially methylated regions);
PHDs: (placenta hypomethylated domains);
PBMC: (peripheral blood mononuclear cells)

## Authors’ contributions

TJJ, DvdB, and ME conceived experimental design. Data acquisition was supervised by TJJ. ZZ and CC implemented the data analysis pipeline and processed all data.

TJJ, SKK, and TL performed secondary data analysis. CG performed a portion of the MCIp experimentation. Figures were designed by TJJ. Manuscript was written by TJJ and ME. All authors discussed the results and commented on the manuscript.

## Acknowledgements

The authors thank Farnaz Ehya, Erin McCarthy, and Helen Tao for performing library preparation and sequencing; Jessica Torres for assistance with figures; Dr. Charles Cantor and Dr. Ron Lindsay for critical review of the manuscript; and Dr. Allan Bombard and Graham McLennan for sample acquisition.

## Additional files

Additional file 1–Supplemental Figure 2

## Additional file 2 – Additional Supplemental Information

This file contains all supplemental figures referred to within the text of the manuscript.

